# Generation of stress fibers through myosin-driven re-organization of the actin cortex

**DOI:** 10.1101/2020.06.30.179283

**Authors:** JI Lehtimäki, EK Rajakylä, S Tojkander, P Lappalainen

## Abstract

Contractile actomyosin bundles, stress fibers, govern key cellular processes including migration, adhesion, and mechanosensing. Stress fibers are thus critical for developmental morphogenesis. The most prominent actomyosin bundles, ventral stress fibers, are generated through coalescence of pre-existing stress fiber precursors. However, whether stress fibers can assemble through other mechanisms has remained elusive. We report that stress fibers can also form without requirement of pre-existing actomyosin bundles. These structures, which we named cortical stress fibers, are embedded in the cell cortex and assemble preferentially underneath the nucleus. In this process, non-muscle myosin II pulses orchestrate the reorganization of cortical actin meshwork into regular bundles, which promote reinforcement of nascent focal adhesions, and subsequent stabilization of the cortical stress fibers. These results identify a new mechanism by which stress fibers can be generated *de novo* from the actin cortex, and establish role for stochastic myosin pulses in the assembly of functional actomyosin bundles.

## Introduction

Cell migration, morphogenesis, and adhesion depend on contractile networks composed of actin and non-myosin II (NMII) filaments. The forces in these structures are generated through sliding of bipolar NMII filaments along actin filaments. Depending on the type of NMII isoform as well as on the organization and dynamics of actin filaments, different types of functional contractile actomyosin arrays can be generated in eukaryotic cells (Lehtimäki et al., 2016). Assembly and contractility of these actomyosin arrays are controlled by upstream signals including Rho-family GTPases, kinase-phosphatase pathways, and Ca^2+^ influxes (Burridge and Guilluy, 2016; Prager-Khoutorsky et al., 2011; Tojkander et al., 2015, 2018; Vicente-Manzanares et al., 2009).

Beneath the plasma membrane of animal cells lies a thin meshwork that is composed of actin filaments, NMII filaments, and associated proteins. This structure, the actin cortex, contributes to morphogenesis of interphase cells, and drives cell rounding during cytokinesis (Bray and White, 1988; Chugh and Paluch, 2018). In human cells, the assembly of actin cortex depends on both formins and the Arp2/3 complex (Bovellan et al., 2014; Lu et al., 2017). Ezrin, radixin, moesin (ERM)-family proteins link the cortical actin meshwork to the plasma membrane in interphase cells and during mitosis (Bretscher et al., 2002). Cortical actin meshwork is constantly under isotropic tension due to intracellular hydrostatic pressure and NMII –generated contractile forces to the actin cortex, and this can lead to formation of membrane blebs through local detachment of the actin cortex from the plasma membrane (Charras and Paluch, 2008). In a three-dimensional environment, some cell types can polarize and persistently migrate by stabilizing the bleb expansion through rearward cortical flows and by modulating the cortical tension (Logue et al., 2015; Ruprecht et al., 2015).

Whereas the actin cortex is composed of an irregular meshwork of actin filaments, animal cells harbor also more highly-ordered actomyosin structures. In many interphase cells, the most prominent actomyosin structures are thick bundles, called stress fibers. These contractile actin bundles often connect to focal adhesions at their ends, and they are especially prominent in cells plated on stiff matrix or when external force is applied to the cells. Thus, stress fibers, together with focal adhesions, constitute a major mechanosensitive machinery in cells (Livne and Geiger, 2016; Tojkander et al., 2012). Apart from adhesion and mechanosensing, stress fibers contribute to cell morphogenesis and tail retraction during migration. Stress fibers also serve as precursors for sarcomeres during cardiomyocyte myofibrillogenesis (Fenix et al., 2018), and stress fiber–like actomyosin bundles contribute to interactions of epithelial cells with their neighbors and with basal lamina (Munjal et al., 2015; Rajakylä et al., 2020; Yamada and Nelson, 2007).

Stress fibers can be classified into different sub-types based on their protein compositions and association with focal adhesions. *Ventral stress fibers* are thick actomyosin bundles that are connected from their both ends to focal adhesions at the bottom of the cell. Despite their name, the central regions of ventral stress fibers often rise towards the dorsal surface of cell (Burnette et al., 2014; Naumanen et al., 2008). In many cell-types, ventral stress fibers associate with each other to form a complex, mechanically interconnected network (Kassianidou et al., 2017; Xu et al., 2012). *Dorsal stress fibers* (also known as radial stress fibers) are non-contractile actin filament bundles that are generated at the cell front through formin mediated actin filament assembly at focal adhesions (Hotulainen and Lappalainen, 2006; Tee et al., 2015). *Transverse arcs*, on the other hand, are thin, contractile actomyosin bundles, arising through NMII-promoted condensing of the lamellipodial actin network at the cell edge (Burnette et al., 2011; Hotulainen and Lappalainen, 2006; Tojkander et al., 2011). Following their appearance, transverse arcs undergo retrograde flow towards the cell center, fuse with each other into thicker bundles, and become more contractile due to concatenation and persistent expansion of NMII filaments into stacks (Beach et al., 2017; Fenix et al., 2016; Hu et al., 2017; Jiu et al., 2019; Tojkander et al., 2015). Although transverse arcs are not directly linked to focal adhesions, they associate with focal adhesion –connected dorsal stress fibers (Burnette et al., 2014). Ventral stress fibers can be generated from the transverse arc and dorsal stress fiber network through a complex process that involves both mechanosensitive regulation of actin filament assembly at focal adhesions as well as inhibition of actin disassembly within the stress fiber network (Hayakawa et al., 2011; Lee and Kumar, 2020; Tojkander et al., 2015, 2018). Moreover, pre-existing ventral stress fibers can undergo ‘splitting’ to generate new, adjacent ventral stress fibers (Young and Higgs, 2018). However, whether stress fibers can also be generated though other mechanisms has remained elusive.

Here we show that different cell-types also exhibit, at their ventral actin cortex, thin stress fibers that are connected to focal adhesions at both ends. These actomyosin bundles, which we named cortical stress fibers, form predominantly underneath the nucleus, and are less contractile and more dynamic compared to the ventral stress fibers, which are derived through fusion of transverse arcs. Importantly, we demonstrate that cortical stress fiber assembly does not involve transverse arcs or any other stress fiber precursor, but that they are generated *de novo* from the actin cortex through NMIIA-driven re-organization of the actin filament meshwork.

## Results

### The actin cortex harbors cortical stress fibers of various size and orientation

Ventral stress fibers were originally defined as contractile actomyosin bundles, which attach to focal adhesions at their both ends (Hotulainen and Lappalainen, 2006; Small et al., 1998). However, migrating mesenchymal cells harbor ventral stress fibers of various length, orientation and thickness, and these can either locate entirely at the ventral surface of cells or rise towards the dorsal surface from their central regions (Baird et al., 2017; Burnette et al., 2014; Elkhatib et al., 2014; Kumari et al., 2020; Lehtimäki et al., 2017; Prager-Khoutorsky et al., 2011; Schulze et al., 2014). To uncover the possible molecular differences between these diverse stress fibers, we utilized the 3D-structured illumination microscopy (SIM) on human osteosarcoma (U2OS) and mouse embryonic fibroblast (MEF) cells migrating on fibronectin. Consistent with previous literature, NMIIA containing, focal adhesion-attached stress fibers of varying thickness and length were visible in both cell lines (Fig. 1A, white and red arrows). In addition to thick ventral stress fibers that connect focal adhesions located at the opposite sides of the cell, both cell-types exhibited thin and relatively short actomyosin bundles that were connected to small focal adhesions at their both ends. As illustrated by the temporal-color coded 3D-SIM projections of F-actin, these thin actomyosin bundles reside at the immediate vicinity of the ventral cortex of the cell (Fig. S1A, white arrows), whereas typical ventral stress fibers (Burnette et al., 2014; Tojkander et al., 2015) rise towards the dorsal surface from the middle of the bundle (Fig. 1A, Fig. S1A; red arrows). Thus, we named these thin, actin cortex – associated actomyosin bundles as *cortical stress fibers*.

**Figure 1.**
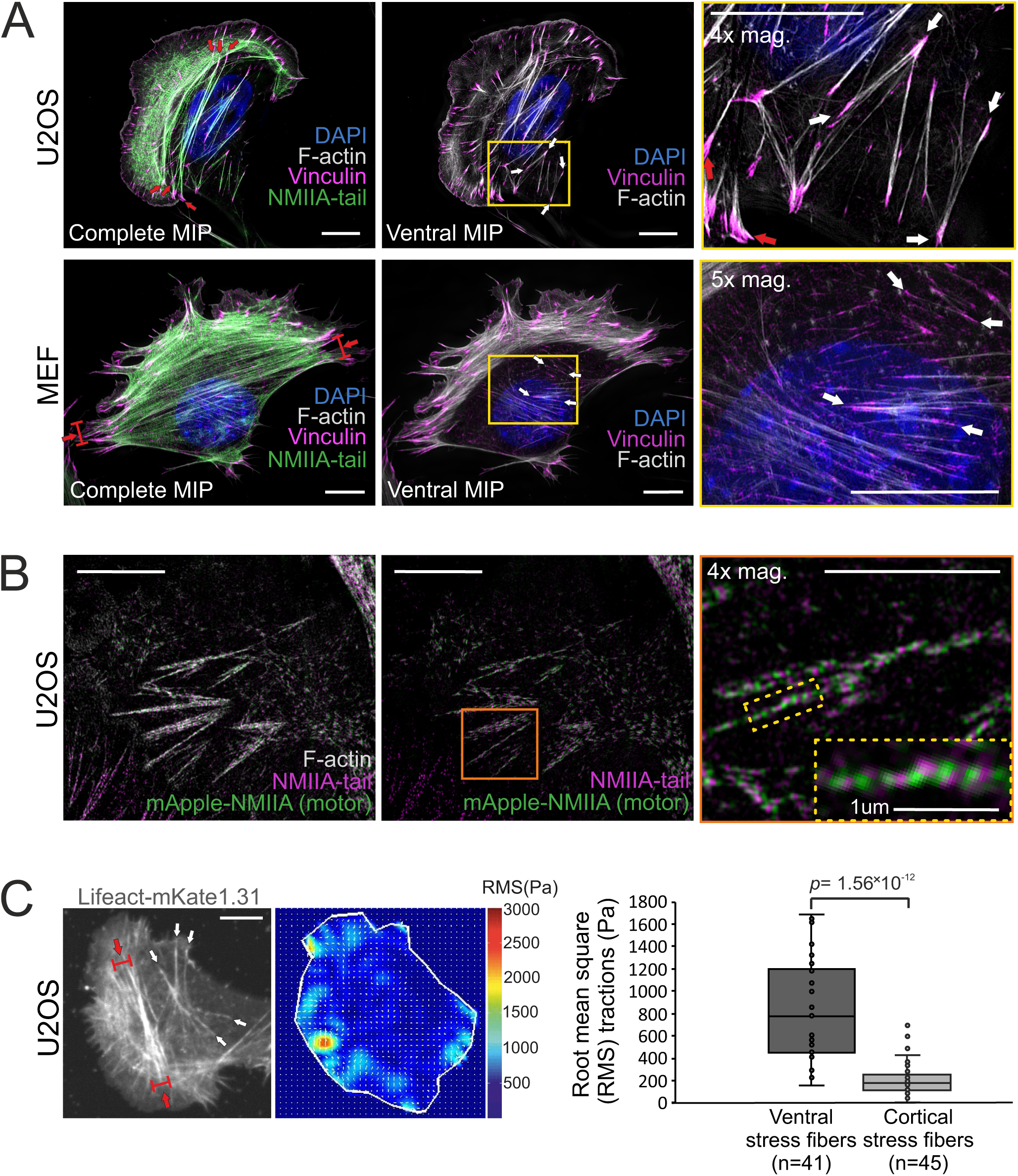
Stress fiber architecture of migrating cells. 3D-SIM maximal intensity projections (MIPs) of the actomyosin networks in cells migrating on fibronectin. **A)** U2OS and mouse embryonic fibroblast (MEF) cells, where the panels on the left display complete MIPs. The panels in the middle show only the filament structures close to the ventral plane (‘ventral MIP’), and the panels on the right are magnifications of the boxed regions from the middle panels. Red arrows highlight ventral stress fibers, and white arrows indicate examples of thin cortical stress fibers that are embedded at the cell cortex. DAPI (blue) and phalloidin (grey) were applied to mark the F-actin and nucleus, respectively. Vinculin (magenta) and NMIIA tails (green) were detected with respective specific antibodies. **B)** 3D-SIM MIP projection from the ventral plane of a U2OS cell transfected with mApple-NMIIA construct (motor, green) and stained with NMIIA-tail specific antibody (magenta, tail) and phalloidin to visualize F-actin. 4x and 10x magnifications (orange box and yellow dotted box, respectively), show the bipolar NMIIA filaments in cortical stress fibers. **C)** Traction force microscopy (TFM) analysis of the forces exerted by ventral stress fibers (red arrows) and cortical stress fibers (white arrows) to the underlying substrate. On the left, exemplary image of a LifeAct-mKate1.31-expressing U2OS cell and the obtained force map with root mean square tractions (RMS). Quantification of the RMS forces between the two stress fiber subtypes (n= 41 for ventral stress fibers and 45 for cortical stress fibers) is shown as box plot on the right. *p*= 1.56 × 10^−12^ (Mann-Whitney *U* test). Scale bars 10 μm and 5 μm for whole cell images and magnifications, respectively.

Similarly to the ventral stress fibers, NMII filaments displayed a bipolar arrangement in the small, basally located cortical stress fibers (Fig. 1B). However, NMII did not appear to assemble into stacks, and the periodic pattern of actin filament cross-linking protein α-actinin was less regular in cortical stress fibers compared to transverse arcs and ventral stress fibers (Fig. S1B). Moreover, the forces exerted by cortical stress fibers to the ECM were much smaller compared to typical ventral stress fibers (Fig. 1C). Although cortical stress fibers were able to form also on laminin and collagen, they were most prevalently observed in cells migrating on fibronectin (Fig.1 and Fig. S1C). Due to their apparent fibronectin preference, we investigated the cortical stress fibers for fibrillar fibronectin deposits and Y118-phospho(p)-paxillin, two markers that are typical to fibrillar and focal adhesions, respectively (Geiger and Yamada, 2011; Zaidel-Bar et al., 2007). Apart from few adhesions in MEFs, the ends of cortical stress fibers did not associate with fibronectin deposits (Fig. S2A, white arrows). Moreover, the adhesions at the ends of cortical stress fibers contained Y118-phospho(p)-paxillin, indicating that they were not associated with fibrillar adhesions (Fig. S2A-B, white arrows). Taken together, the molecular composition of cortical stress fibers closely resembles the one of ventral stress fibers, but they are typically much smaller, localize consistently at the ventral actin cortex, and exert only weak traction forces.

### Cortical stress fibers are generated *de novo* from the ventral actin cortex

Ventral stress fibers are generated from a network of pre-existing transverse arcs and focal adhesion-attached dorsal stress fibers (Tojkander et al., 2015, 2018). Thus, we examined if the thin cortical stress fibers are generated by the same or a different mechanism. To this end, we imaged the ventral region of migrating U2OS cells and MEFs expressing LifeAct-TagGFP2 (to detect F-actin) and vinculin-mApple (to visualize focal adhesions) by time-lapse total internal reflection microscopy (TIRFM). Surprisingly, these experiments revealed that that cortical stress fibers emerged *de novo* from the ventral actin cortex, without involvement of any pre-existing stress fiber precursors (Fig. 2A-B and Movie S1). In this process, actin filaments of the cell cortex re-organized into thicker bundles (Fig. 2A-B, blue arrows) and this was followed by growth of nascent, initially barely visible, vinculin-positive adhesions at the both ends of the bundle (Fig. 2A-B, orange arrows). This eventually led to a formation of an actin filament bundle that was connected to focal adhesions at its both ends.

**Figure 2.**
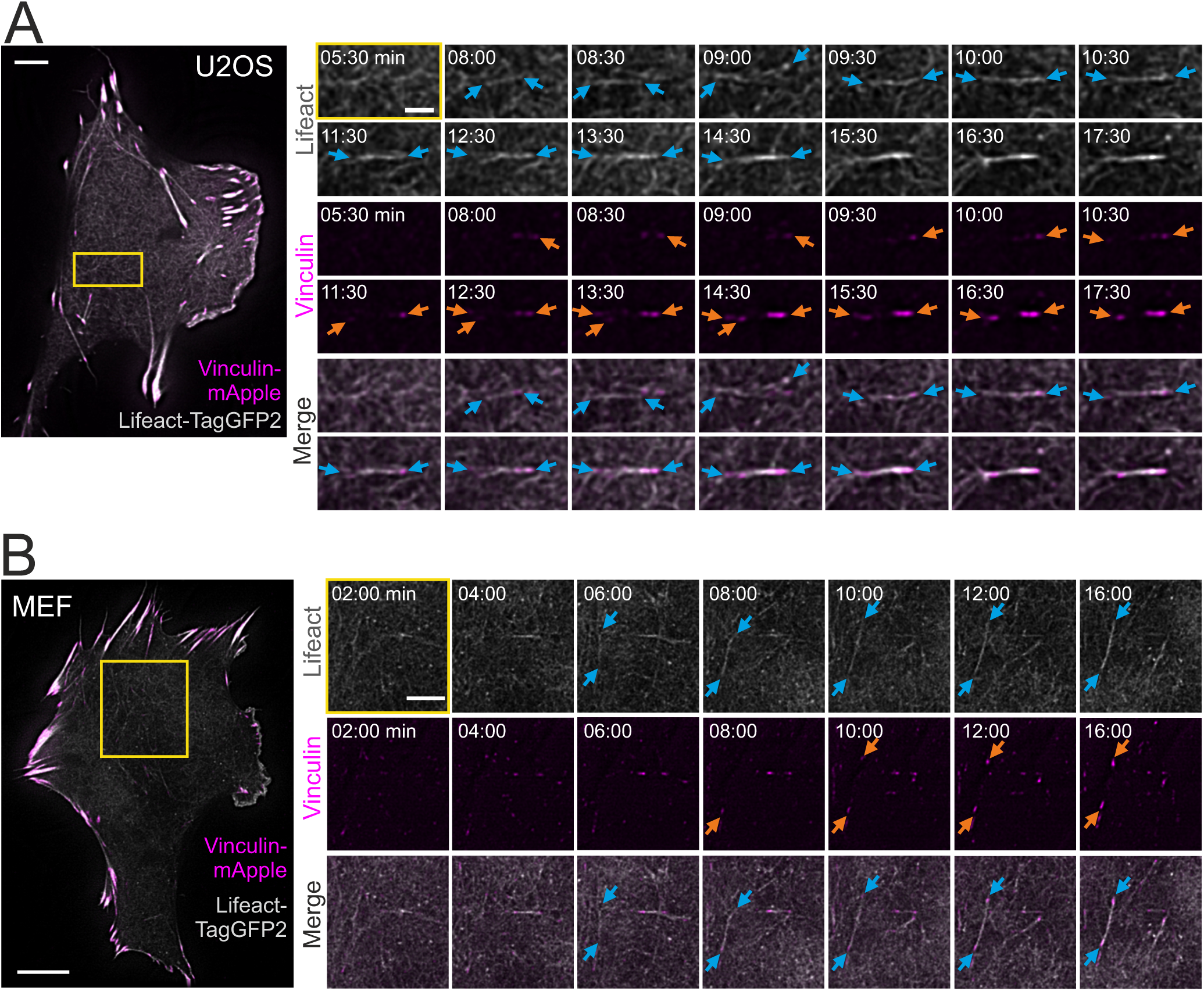
Cortical stress fibers assemble *de novo* from the actin cortex. TIRF time-lapse imaging of a migrating U2OS **(A)** and MEF cell **(B)**, expressing LifeAct-TagGFP2 (grey) and vinculin-mApple (magenta). Selected time-lapse frames from the magnified area (yellow box) are shown on the right as separate channels and merged frames. These demonstrate the *de novo* emergence of a cortical stress fiber from the ventral actin cortex. Blue arrows illustrate F-actin bundling and orange arrows point the maturation of the vinculin-positive focal adhesions. See also **Movie S1**. Scale bars 10 μm and 5 μm for whole cell images and time-lapse zoom-ins, respectively. Imaging interval 30s.

Live-cell imaging experiments also demonstrated that cortical stress fibers are typically very dynamic with relatively short half-life, they often associate with each other, and can arise via various intermediate assembly states (Fig. S2C and Movie S2). Collectively, these data reveal that cortical stress fibers assemble through a novel mechanism from actin cortex, without involvement of stress fiber precursors.

### Cortical stress fibers preferably emerge underneath the nucleus in migrating cells

Cortical stress fibers were enriched at the rear of migrating cells, and were typically located either underneath or close to the nucleus (Fig. 1). Nucleus is a relatively bulky organelle, localized behind the lamella at the cell rear in polarized mesenchymal cells, and it undergoes NMII-dependent translocation as the cell moves forward (Thomas et al., 2015; Wu et al., 2014). Thus, we examined possible connection between the nucleus and generation of cortical stress fibers. Live-imaging of U2OS cells expressing Histone-H2B-mCherry to mark the nucleus, as well as LifeAct-TagGFP2 and focal adhesion marker miRFP670-paxillin, revealed that the assembly of cortical stress fibers occurred typically under the nucleus during cell locomotion (Fig. 3A and Movie S3). Similar results were obtained when nuclei were visualized together with miRFP670-paxillin and mApple-NMIIA (Fig. S3A). Quantification of the emergence of cortical stress fibers from 19 time-lapse movies revealed that ∼80 % of the cortical stress fibers assembled underneath the nucleus (Fig. 3B). By considering that the nucleus typically consist <30 % of the cell area, the assembly of cortical stress fibers occurs predominantly underneath the nucleus.

**Figure 3.**
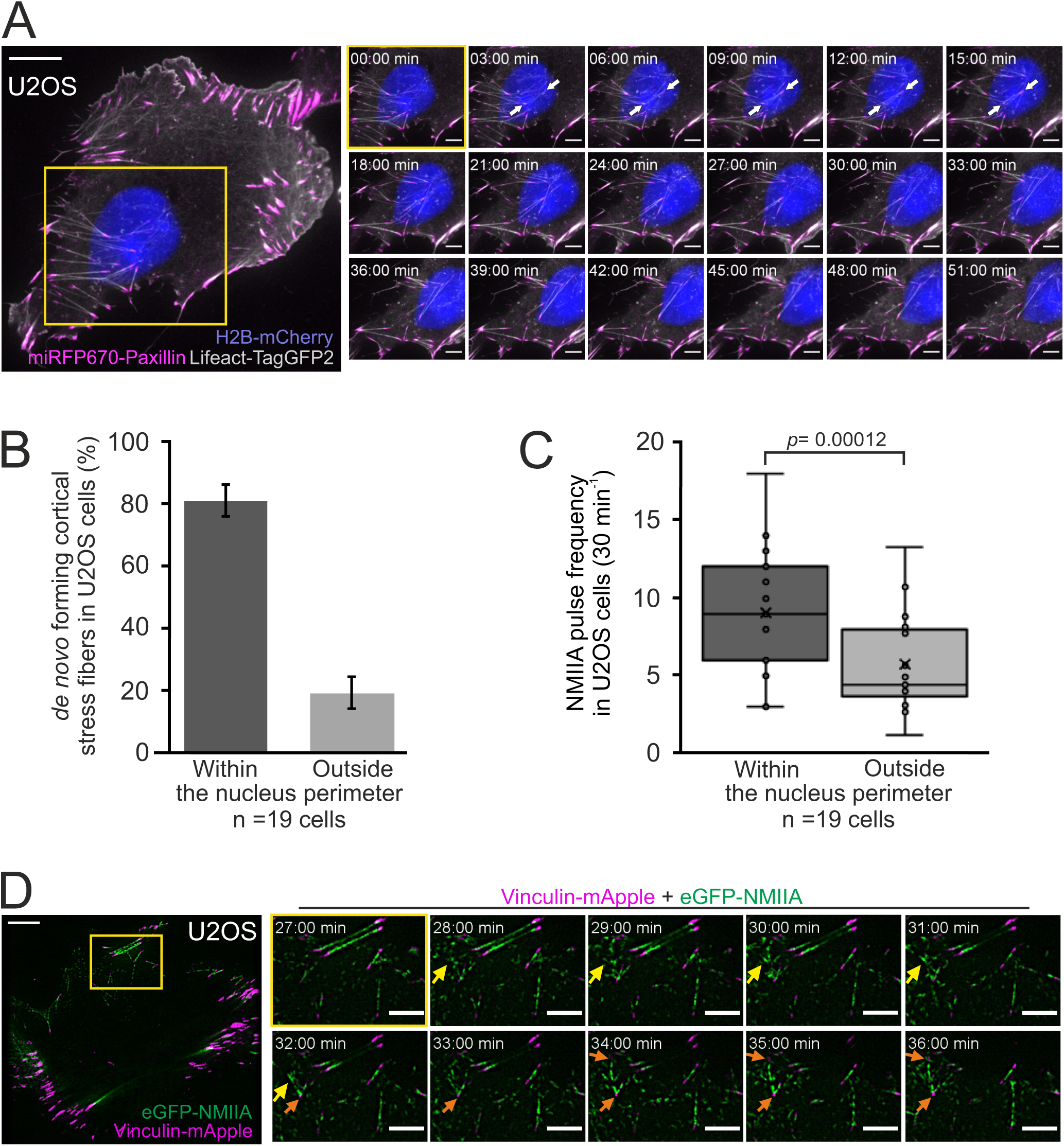
Cortical stress fibers assemble predominantly underneath the nucleus in migrating cells. **A)** TIRF time-lapse imaging of a migrating U2OS cell. The selected time-lapse frames (from the yellow boxed area) illustrate the assembly of a cortical stress fiber (exemplified by white arrows) below the moving nucleus. Histone H2B-mCherry was applied to detect the nucleus (blue, EPI-TIRF) and miRFP670-paxillin (magenta) and LifeAct-TagGFP2 (grey) to visualize focal adhesions and F-actin, respectively. See also **Movie S3. B)** Quantification of the position of *de novo* formation of cortical stress fibers from TIRF-time-lapse videos of U2OS cells expressing LifeAct-TagGFP2, Histone-H2B-mCherry, and miRFP670-paxillin. Data are presented as %-distribution of cortical stress fibers assembled under the nucleus versus outside the nucleus perimeter ± SEM. n = 101 cortical stress fibers analyzed from 19 cells. **C)** Quantification of NMIIA pulse location at the ventral cortex of U2OS cells expressing Histone H2B-eGFP and mApple-NMIIA. Data are presented as pulse frequency (number of individual pulses) under the nucleus vs. outside the nucleus perimeter (rest of the ventral cortex). Each data point is normalized to the area of nucleus. Box plot displays median, outlier whiskers and ‘x’ points out the mean. n= 19 cells. Significance (*p* = 0.00012) tested with a paired *t*-test. **D)** TIRF time-lapse imaging of a migrating U2OS cell expressing eGFP-NMIIA (green) and vinculin-mApple (magenta). Selected time-lapse frames from the magnified area (yellow box) demonstrate that NMIIA pulse (yellow arrow) is associated with the assembly of cortical stress fibers and enforcement of vinculin positive focal adhesions (orange arrows). See also **Movie S2**. Scale bars 10 μm and 5 μm for whole cell images and time-lapse zoom-ins, respectively. Imaging interval 30s.

### Myosin pulses direct cortical stress fiber assembly and maturation

Pulsatile behavior of NMII at the actin cortex has been documented in epithelial and mesenchymal cells, and reported to induce transient accumulation of cortical actin (Baird et al., 2017; Kim and Davidson, 2011; Munjal et al., 2015). Our analysis indicated that NMIIA pulses at the ventral cortex of U2OS cells are slightly more prevalent underneath the nucleus where cortical stress fibers also principally assembled, compared to the region outside the nucleus perimeter (Fig. 3C). To reveal the possible role of NMII pulses in generation of the cortical stress fibers, we performed TIRFM imaging of U2OS cells expressing eGFP-NMIIA and vinculin-mApple. Interestingly, assembly of cortical stress fibers, and enforcement of focal adhesions at their ends, were associated with NMIIA pulses (Fig. 3D and Movie S2). To examine this in more detail, we performed three-color TIRFM on U2OS cells expressing mApple-NMIIA, LifeAct-TagGFP2, and miRFP670-paxillin. These experiments revealed that NMIIA pulses frequently coincided with transient F-actin bundling (Fig 4A and Movie S4). The forming actomyosin bundle engaged paxillin positive focal adhesions, causing them to align to the direction of the pull and to increase in size, thus leading to stabilization and maturation of the cortical stress fiber. Importantly, focal adhesions associating with the new cortical stress fiber could either emerge *de novo* (Fig. S3B, orange arrow, Movie S5), or be pre-existing adhesions that were connected to other, discrete contractile bundles (Fig. S2C, Fig. 4B, Fig. S3C and Movie S4-S5). Moreover, we occasionally observed assembly of cortical stress fibers, where one end of the contractile bundle was connected to the pre-existing actomyosin bundles instead of a focal adhesion (Fig. S3B, yellow arrow, and Movie S5). Thus, generation of cortical stress fibers is both very dynamic and plastic process, and it can involve either *de novo* formation of focal adhesions, or ‘re-cycling’ of pre-existing focal adhesions or higher-order actomyosin structures to connect the nascent cortical stress fiber to the cell cortex.

**Figure 4.**
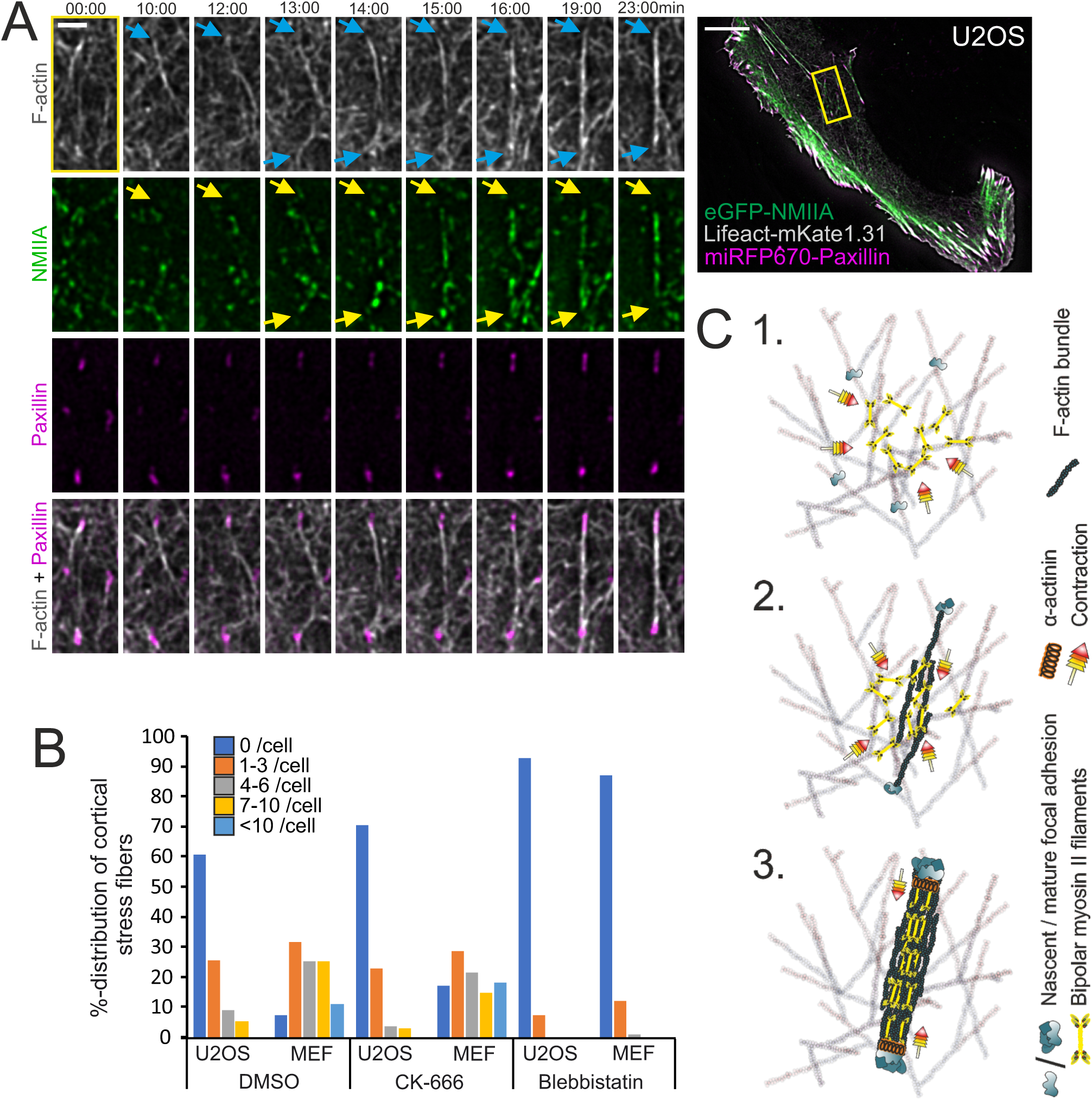
Cortical NMII pulses orchestrate F-actin bundling for the assembly of cortical stress fibers. **A)** Formation of a cortical stress fiber from the ventral cortex of a migrating U2OS cell as studied by time-lapse TIRF microscopy. The entire cell is shown on right, and the selected time-lapse frames (from the region indicated by yellow box) display how F-actin (grey, LifeAct-mKate2; blue arrows) and NMIIA (green, eGFP-NMIIA; yellow arrows) organize into an actomyosin bundle, which promotes the enlargement of pre-existing paxillin-positive focal adhesions (FA) (magenta, miRFP670-paxillin). Note that prior to bundle assembly, the pre-existing focal adhesions were connected to different actomyosin bundles. Original imaging interval 30s. See also **Movie S4**. Scale bars 10 μm and 5 μm for whole cell image and magnified time-lapse frames, respectively. **B)** Percentual distribution of cortical stress fiber numbers in U2OS cells and MEFs after different pharmacological perturbations. Cortical stress fiber number and length measurements were obtained through blind analysis of the tile-scan TIRF data. n= DMSO 157 and 247, CK-666, 166 and 182, Blebbistatin 195 and 198 cells for U2OS cells and MEFs, respectively. See also **Figure S4A** and **S4C. C)** Schematic representation of the *de novo* cortical stress fiber assembly from the ventral actomyosin cortex. 1. NMIIA pulses occur frequently at the ventral actin cortex. 2. These pulses can cause transient accumulation and bundling of the cortical actin filaments via myosin-mediated actin filament crosslinking and re-organization. This triggers the enlargement of nascent focal adhesions at the ends of the bundle. 3. The transient actomyosin bundles can mature to a cortical stress fiber through recruitment of more actin filaments and NMII.

To examine the role of NMII in the assembly of cortical stress fibers more closely, we applied NMII inhibitor para-amino-Blebbistatin. As reported earlier (Baird et al., 2017), Blebbistatin did not cease the NMII pulsatile behavior in U2OS cells. However, the cortical stress fiber formation was almost completely abolished, providing evidence that NMII activity is critical for the assembly of cortical stress fibers (Fig. S4A, and Movie S6). We also cultured U2OS cells on poly-D-lysine coated imaging dishes to examine the importance of integrin-based adhesions in this process. By imaging cells that had managed to adhere to the dish only from their edges, but were devoid of adhesions inside cell margins, we revealed that while NMII pulses still drove transient cortical F-actin enrichment, they could not engage F-actin bundling to the assembly of cortical stress fibers (Fig S4B, and Movie S7). Finally, we applied the Arp2/3 inhibitor CK-666 to elucidate the possible role of Arp2/3 complex mediated actin filament assembly in the generation of cortical stress fibers. Even after 4 hours post drug administration, *de novo* cortical stress fiber assembly was frequently observed (Fig. S4C, and Movie S8). Quantification of cortical stress fibers from control (DMSO-treated), para-amino-Blebbistatin, and CK666 treated U2OS cells and MEFs confirmed that NMII inhibition with this compound led to an almost complete loss of cortical stress fibers, whereas Arp2/3 inhibition resulted in only a moderate decrease in the amount of these actomyosin bundles (Fig 4B). Collectively, these results demonstrate that generation of cortical stress fibers is driven by NMII-catalyzed re-organization of the cortical actin filament network (Fig. 4C).

## Discussion

Here we discovered a novel mechanism by which focal adhesion –attached stress fibers assemble in cells. We show that unlike ventral stress fibers, which are generated from the pre-exiting network of dorsal stress fibers and transverse arcs (Tojkander et al., 2015), cortical stress fibers located at the ventral cortex of 2D-migrating assemble *de novo* through NMII-dependent re-organization of the actin cortex. Cortical stress fibers bear close resemblance to the ventral stress fibers because they terminate to focal adhesions at both ends and are composed of bipolar NMII filaments and actinin decorated actin filaments. However, they are thinner, more dynamic, devoid of NMII stacks and less contractile compared to ventral stress fibers.

The assembly of cortical stress fibers is intimately linked to the actin cortex, where stochastic, NMII-mediated contractions of the cortical actin meshwork can lead to the formation of cortical actomyosin bundles. These are initially transient by nature, and can either form a cortical stress fiber of collapse back into the ventral cortex. Although NMII-pulses have not been previously linked to assembly of contractile actomyosin bundles, transient NMII pulse-driven cortical actin condensations were reported to occur in epithelial tissues of developing organisms where they drive tissue morphogenesis (Michaux et al., 2018; Munjal et al., 2015; Munro et al., 2004; Rauzi et al., 2010; Solon et al., 2009) and in cells of mesenchymal origin migrating in 2D environment (Baird et al., 2017; Graessl et al., 2017; Kim and Davidson, 2011). In *Drosophila* epithelia, the NMII/actin pulses were also reported to undergo E-cadherin-mediated stabilization into medioapical filaments (Mason et al., 2013; Vasquez et al., 2014). Thus, myosin pulse-driven F-actin bundling may have a more general role in formation of a different types of contractile actomyosin bundles in cells.

Also during cytokinesis, cortical actin flows drive the assembly of small actomyosin nodes with random F-actin orientation. These become organized as elongated bundles with some resemblance to stress fibers (Henson et al., 2017; Laplante et al., 2016). Thus, initially very similar, nascent cortical actomyosin foci appear drive the formation of various different contractile structures. In line with stochasticity of the myosin pulses, also the nascent cortical stress fibers come in many intermediate shapes that reflect the initially random formation of NMII/F-actin clusters. These intermediate structures can become momentarily engaged by more than two focal adhesions, but during subsequent tug-of-war-type process usually two focal adhesions become dominant at the ends of the cortical stress fiber. The degree of cortical F-actin network bundling may be dependent on both the mesh gap-size and thickness of the actin meshwork that vary between different cell types (Bovellan et al., 2014; Svitkina, 2020). Interestingly, the, actin cortex appears to be more dense and under higher tension at the back of the 2D-migrating cells (Bisaria et al., 2020; Chugh and Paluch, 2018), where we observe the most prominent cortical stress fiber assembly.

What could be the reason underlying the assembly of cortical stress fibers under the nucleus in migrating mesenchymal cells? Transportation of the relatively rigid nucleus in cells migrating in tissue environment poses mechanical challenges, especially when facing narrow passages (Denais et al., 2016; Petrie et al., 2014). This results in accumulation of contractile F-actin bundles around the nucleus to mediate movement of the nucleus through the constriction (Davidson et al., 2020; Thomas et al., 2015). Similarly, when subjecting MEF cells to adhere and migrate along narrow ECM-coated tracks, directional F-actin bundles are observed under the nucleus. The stress fibers underneath the nucleus appear to be largely independent of linker of nucleus and cytoskeleton (LINC)-complex, which is known to mediate contacts between the nuclear envelope and perinuclear cap fibers at the dorsal surface of the nucleus (Calero-Cuenca et al., 2018; Kim et al., 2014, 2012; Luxton et al., 2010). Although the mechanisms by which these actomyosin bundles protecting the nucleus are assembled is still unknown, we envision that it would be difficult to generate such fibers from the transverse arc network, which to our knowledge is not present in cell migrating in a 3D environment (Doyle et al., 2015; Thomas et al., 2015). Thus, the *de novo* mechanism to generate contractile cortical stress fibers reported here is well-suited for generation of stress fibers in 3D environment to protect the nucleus during migration. In line with this hypothesis, cells in 3D environment utilizing mesenchymal migration mode exhibit multiple focal adhesion-mediated ECM attachments around their dorso-ventral and antero-posterior axis, and impose greater challenges with protecting the integrity of the nucleus than cells in 2D (Yamada and Sixt, 2019).

Taken together, we demonstrate that the myosin II-pulse-induced actin contractions are linked to maturation of nascent focal adhesions at the cell cortex, and this can lead to an assembly of a contractile stress fiber that is attached to a focal adhesions at its both ends. This new mechanism expands the toolbox that cells can apply for generation of stress fibers, thus allowing assembly of stress fibers also in conditions where the pre-existing stress fiber precursors (transverse arcs and dorsal stress fibers) are not present. In the future, it will be important to examine the functional relevance of cortical stress fibers especially in 3D environment, where transverse arcs are not present and focal adhesion formation is not restricted to the ventral plane.

## Materials and methods

### Cell culture and transfections

Wild-type mouse embryonic fibroblasts (MEF) (a kind gift from J. Eriksson’s lab) and human osteosarcoma (U2OS) (ATCC HTB-96) cells were cultured in high-glucose DMEM (Lonza, BE12-119F), supplemented with 10% FBS (Gibco, 10500-064), 10 U/ml penicillin, 10 µg/ml streptomycin and 20 mM L-glutamine (Gibco, 10378-016) and kept at +37°C in a humified atmosphere with 5% CO_2_. For live-cell imaging experiments, Fluorobrite DMEM (Gibco, A1896701), supplemented with 25 mM HEPES and 10% FBS was used. Transient transfections were performed with Lipofectamine 2000 (Thermo Fisher Scientific) according to the manufacturer’s instructions. 24 hours after transfection, cells were either fixed with 4% PFA in PBS for 10 min at +37°C (3D-SIM/tile-scan TIRF) or detached with 0.05% Trypsin-EDTA (Gibco, 15400054) and plated onto high-precision (#1.5H) 35 mm imaging dishes (Ibidi μ-dish high, 81158), coated with 10 µg/ml fibronectin (Merck, FIBRP-RO) for 1 h RT and placed +4° C overnight prior usage in live-cell TIRF imaging experiments. To prevent integrin-mediated adhesion, 10 µg/ml poly-D-lysine (Merck, P0899) was applied onto the 35 mm imaging dishes at RT for 10 minutes, washed, and allowed to dry prior plating the cells. Cells were allowed to attach for 24 h prior onset of imaging. For 3D-SIM or tile-scan TIRF, cells were plated onto #1.5H coverslips coated with 10 μg/ml fibronectin (Merck, FIBRP-RO), 25 μg/ml laminin (Merck, L2020), or 500 μg/ml collagen (Merck, C4243).

### Plasmids

eGFP-NMIIA was a gift from Mathew Krummel (Addgene plasmid #38297; Jacobelli et al., 2009). LifeAct-mKate1.31, mApple-NMIIA, Vinculin-mApple and α-actinin1-TagRFP-T were all gifts from Michael Davidson (Addgene plasmids #54668, #54929, #54962, #58033, respectively; Burnette et al., 2014; Shcherbo et al., 2009). LifeAct-TagGFP2 was a kindly provided by the Emmanuel Lemichez lab. Histone H2B-eGFP (in peGFP-N3 backbone) and Histone H2B-mCherry (in pmCherry-N3 backbone) were gifts from Maria Vartiainen lab. miRFP670-paxillin (pPL1514) was constructed by replacing the GFP in eGFP-paxillin (containing *gallus gallus* paxillin ORF with T132S and M133L unattended mutations in an eGFP-C1 backbone) with miRFP670 originated from miRFP670-N1 (a gift from Vladislav Verkhusha, Addgene plasmid #79987; Shcherbakova et al., 2016) via restriction free cloning using NEBuilder (New England Biolabs). All inserts have been sequence verified prior the respective plasmid usage.

### Reagents

Mouse monoclonal antibody for vinculin (Merck, V9131), rabbit polyclonal antibody recognizing the C-terminal tail of the NMIIA heavy chain (Biolegend, 909801), rabbit polyclonal anti-fibronectin (F3648, Sigma Aldrich), and monoclonal rabbit antibody against T118-phosphorylated paxillin (#2541, Cell Signaling Technology) were employed in this study. 4’,6’-diamidino-2-phenylindole (DAPI) (Thermo Fisher Scientific, D1306) was utilized to detect the nuclei, whereas Alexa Fluor 488- and 568-Phalloidin (Thermo Fisher Scientific, A12379/12380) were applied to visualize F-actin. Alexa Fluor 488- and 568-congujated goat anti-mouse (Thermo Fisher Scientific, A-11001 and A-11031, respectively) and Alexa Fluor Plus 647-conjugated goat anti-rabbit (Thermo Fisher Scientific, A32733) were used as secondary antibodies. 50 μM of para-amino-Blebbistatin (Optopharma, DR-Am-89) was applied for one hour to inhibit NMII activity and 100 μM CK-666 (Merck, SML0006) for four hours to inhibit the Arp2/3 complex. Both inhibitor stocks were made to dimethyl sulfoxide (DMSO) (Merck, D2650) that was also used as control in the corresponding experiments. For 3D-SIM experiments, samples were mounted using either non-hardening Vectashield (Vector laboratories, H-1000) or Prolong Glass (Thermo Fisher Scientific, P36980).

### 3D-Structured Illumination Microscopy (SIM)

All 3D-SIM images were obtained at RT, using Deltavision OMX SR (Cytiva) with 60x/1.42NA PlanApo N oil objective with 1.516 RI immersion oil, a laser module with 405-, 488-, 568- and 640 nm diode laser lines and three sCMOS cameras, operated through Acquire SR 4.4 acquisition software. SI reconstruction and image alignment were performed with SoftWoRx 7.0. Imaging arrays of 1024 × 1024 or 512 × 512 were used, both with pixel size of 0.08 µm and 0.125 µm (x/y and z). Samples for 3D-SIM were prepared according to (Kraus et al., 2017) with exceptions of using 5% BSA-PBST as blocking reagent, and omitting the pre-incubation with mounting reagent when using the Prolong Glass as a mountant.

### IRF microscopy

For live-cell TIRF experiments, the Ring-TIRF module from the Deltavision OMX SR (Cytiva) with 60x/1.49NA Apo N oil objective (Olympus), 1.522 RI immersion oil and imaging chamber with controlled humified atmosphere of 5% CO_2_ and 37°C was utilized. Sample illumination with 488, 568 and 640 nm diode lasers was detected and recorded with three respective sCMOS cameras and controlled through Acquire SR 4.4 acquisition software. The captured 1024 × 1024 time-lapse videos had a pixel size of 0.08 µm (x/y). Obtained time-lapse series were deconvolved and channels aligned with SoftWoRx 7.0. Prior the onset of live-cell imaging, cells were allowed to settle within the imaging chamber for 1 hour.

‘Tile-scan’ TIRF of fixed specimens on 35 mm imaging dishes (Ibidi μ-dish high) in 1x PBS, was performed with Eclipse Ti-E N-STOM/TIRF microscope (Nikon Instruments) using 100x/1.49NA Apo TIRF oil objective. Illumination was provided by 405 nm (100mW),488 nm (Argon) and 561 nm (150mW) laser lines and LU4A laser unit (Nikon Instruments), controlled via NIS-Elements (NIS-AR version 4.5). Images were captured using iXon+ 885 EMCCD camera (Andor Technology) with imaging array of 1004 x 1002 pixels, and final pixel size of 0.08 µm (x/y). In order to acquire the data set from the specimen in unbiased manner, focus was set, followed by capturing 5×5 field of view (FOV) tile-scan with 10% overlap. After this, the next tile-scan was acquired from a new area, six times the FOV to another direction.

### Traction force microscopy

For the TFM experiments, U2OS cells on a 35 mm imaging dishes (Mat-Tek, P35G-1.5-14C) were transfected with mKate1.31-LifeAct using JetPRIME transfection reagent (Polyplus transfection, 114-01) according to the manufacturer’s instructions. Next day, cells were re-plated on polyacrylamide (PAA) gels of known stiffness (Young’s Modulus/elastic modulus = 26 kPa) coated with mixture of collagen and fibronectin and incubated for 2-4 h prior to imaging. PAA substrates were surface-coated with sulfate fluorescent microspheres of 0.2 µm diameter (Invitrogen, F8848) before coating with the ECM proteins. Cells and the underlying microspheres were imaged with 3I-Marianas imaging setup containing a heated sample chamber (+37°C) and controlled 5% CO_2_ (3I intelligent Imaging Innovations, Germany). 63x/1.2 W C-Apochromat Corr WD=0.28 M27 objective was used. After first round of images, the cells were removed from the substrates with 10 x Trypsin (Lonza, BE02-007E) and a second set of images were obtained of the microspheres in a cell-free configuration. Microsphere displacement maps were achieved by comparing the first and second set of microsphere images. By knowing the spatial displacement field, substrate stiffness (26 kPa), and tracing manually cell boundaries and single adhesions in the ends of stress fibers, we could compute the traction fields by using Fourier Transform Traction Cytometry (Krishnan et al., 2009; Tolić-Nørrelykke et al., 2002). From the traction fields, root mean squared (RMS) magnitudes were computed. Importantly, several of the cortical stress fibers relayed forces so low that they were undetectable by the microsphere displacement assay, thus the actual RMS-values for this stress fiber subtype are likely to be even lower. Distinct stress fiber types for the measurements were chosen based on their appearance, location and connections to FAs: Ventral stress fibers were defined as thick, straight bundles, usually behind an arc network and connected to focal adhesions from both ends; Cortical stress fibers were defined as thin and short bundles, usually located under the nucleus and associated with focal adhesions from both ends.

### NMIIA pulse quantification

A Fiji/ImageJ macro originally used to quantify F-actin dynamics was obtained from Lance Davidson (Kim and Davidson, 2011), and modified to detect and track the frequency of NMIIA pulses. Pulses were tracked at the ventral cortex from TIRF time-lapse series of U2OS cells expressing mApple-NMIIA and H2B-eGFP, and analyzed similar to (Baird et al., 2017) with exceptions of defining the initial regions of interest (ROIs) so that pulses were recorded first from the ventral cortex excluding nucleus (‘outside’) and then only within the nucleus perimeter (‘within’). Also more stringent 1.3 threshold of NMIIA signal over the background intensity was used and number of connected hexagons smaller than 3 (in individual x-y frames) were filtered out to improve the analysis by excluding single bright myosin puncta. Furthermore, the ‘outside’ ROI excluded edges of the cell as well as the early lamella, to prevent the NMII signal from these locations to bias the analysis. To minimize the effect of photobleaching, pulse frequency was determined by registering all pulses within the first 30 minutes of TIRFM time-lapse videos that persisted more than three frames. The frequency of each ‘outside’ data point was further normalized to the smaller ventral cortex area covered by the nucleus as calculated from the drawn ROIs with Fiji/ImageJ (Schindelin et al., 2012).

### Image processing and statistical analyses

All TIRFM data, excluding the LifeAct channel, was processed with the rolling ball background subtraction plug-in using a 50-pixel ball radius in Fiji/ImageJ. For 3D-SIM data, temporal color-coded hyperstacks were also created with Fiji. The statistical analysis for TFM data, NMII pulse frequency and cortical stress fiber localization, as well as respective figure generation, were performed with Excel (Microsoft). Statistical significance for myosin pulse frequency was calculated with paired *t*-test, preceded by examining data normality with quantile-quantile plot. To assess statistical difference for the TFM data, Mann-Whitney *U*-test was applied. In quantification of cortical stress fiber formation from the TIRFM videos (Fig. 3B), cortical stress fibers of maximum 10 µm diameter, persisting for 2 minutes or longer, having focal adhesion at both ends with possibility of sharing a common focal adhesion, were included in the analysis. To be categorized as forming under the nucleus, at least one focal adhesion had to reside below the nucleus. Number of cells analyzed for each quantification, are listed in the respective figure legends. The cortical stress fibers number form TIRF images of fixed cells treated with Blebbistatin, CK-666 or DMSO alone were quantified by manual blind analysis. Here, the cortical stress fibers were defined as focal adhesion attached (from both ends) actin bundles located at least partially underneath the nucleus and with a length less than the diameter of nucleus.

## Acknowledgements

We would like to thank the Biomedicum Imaging Unit of the University of Helsinki, sponsored by HiLIFE and Biocenter Finland, for support with imaging. Maria Vartiainen (University of Helsinki), Johanna Ivaska (University of Turku) are acknowledged for providing reagents. We thank Lance Davidson, University of Pittsburgh for providing the macro, and Harri Jäälinoja (LMU imaging unit, Institute of Biotechnology, University of Helsinki) for help with modifying the macro for pulse quantification. This work was supported by grants from Sigrid Juselius Foundation (to P.L.) Jane and Aatos Erkko Foundation (to P.L. and S.T.) and Academy of Finland (to S.T.).

## Author contributions

JL and PL designed the study and wrote the paper with comments from all authors. JL conducted all experiments and analyzed the data, expect for the TFM experiments performed and analyzed by ER and ST.

## References

Baird, M.A., Billington, N., Wang, A., Adelstein, R.S., Sellers, J.R., Fischer, R.S., and Waterman, C.M. (2017). Local pulsatile contractions are an intrinsic property of the myosin 2A motor in the cortical cytoskeleton of adherent cells. Mol. Biol. Cell 28, 240–251.

Beach, J.R., Bruun, K.S., Shao, L., Li, D., Swider, Z., Remmert, K., Zhang, Y., Conti, M.A., Adelstein, R.S., Rusan, N.M., et al. (2017). Actin dynamics and competition for myosin monomer govern the sequential amplification of myosin filaments. Nat. Cell Biol. 19, 85–93.

Bisaria, A., Hayer, A., Garbett, D., Cohen, D., and Meyer, T. (2020). Membrane-proximal F-actin restricts local membrane protrusions and directs cell migration. Science (80-.). 368, 1205–1210.

Bovellan, M., Romeo, Y., Biro, M., Boden, A., Chugh, P., Yonis, A., Vaghela, M., Fritzsche, M., Moulding, D., Thorogate, R., et al. (2014). Cellular Control of Cortical Actin Nucleation. Curr. Biol. 24, 1628–1635.

Bray, D., and White, J.G. (1988). Cortical flow in animal cells. Science 239, 883–888.

Bretscher, A., Edwards, K., and Fehon, R.G. (2002). ERM proteins and merlin: integrators at the cell cortex. Nat. Rev. Mol. Cell Biol. 3, 586–599.

Burnette, D.T., Manley, S., Sengupta, P., Sougrat, R., Davidson, M.W., Kachar, B., and Lippincott-Schwartz, J. (2011). A role for actin arcs in the leading-edge advance of migrating cells. Nat. Cell Biol. 13, 371–381.

Burnette, D.T., Shao, L., Ott, C., Pasapera, A.M., Fischer, R.S., Baird, M.A., Der Loughian, C., Delanoe-Ayari, H., Paszek, M.J., Davidson, M.W., et al. (2014). A contractile and counterbalancing adhesion system controls the 3D shape of crawling cells. J. Cell Biol. 205, 83–96.

Burridge, K., and Guilluy, C. (2016). Focal adhesions, stress fibers and mechanical tension. Exp. Cell Res. 343, 14–20.

Calero-Cuenca, F.J., Janota, C.S., and Gomes, E.R. (2018). Dealing with the nucleus during cell migration. Curr. Opin. Cell Biol. 50, 35–41.

Charras, G., and Paluch, E. (2008). Blebs lead the way: how to migrate without lamellipodia. Nat. Rev. Mol. Cell Biol. 9, 730–736.

Chugh, P., and Paluch, E.K. (2018). The actin cortex at a glance. J. Cell Sci. 131.

Davidson, P.M., Battistella, A., Déjardin, T., Betz, T., Plastino, J., Borghi, N., Cadot, B., and Sykes, C. (2020). Nesprin-2 accumulates at the front of the nucleus during confined cell migration. EMBO Rep.

Denais, C.M., Gilbert, R.M., Isermann, P., McGregor, A.L., Te Lindert, M., Weigelin, B., Davidson, P.M., Friedl, P., Wolf, K., and Lammerding, J. (2016). Nuclear envelope rupture and repair during cancer cell migration. Science (80-.). 352, 353–358.

Doyle, A.D., Carvajal, N., Jin, A., Matsumoto, K., and Yamada, K.M. (2015). Local 3D matrix microenvironment regulates cell migration through spatiotemporal dynamics of contractility-dependent adhesions. Nat. Commun. 6, 8720.

Elkhatib, N., Neu, M.B., Zensen, C., Schmoller, K.M., Louvard, D., Bausch, A.R., Betz, T., and Vignjevic, D.M. (2014). Fascin plays a role in stress fiber organization and focal adhesion disassembly. Curr. Biol. 24, 1492–1499.

Fenix, A.M., Taneja, N., Buttler, C.A., Lewis, J., Van Engelenburg, S.B., Ohi, R., and Burnette, D.T. (2016). Expansion and concatenation of nonmuscle myosin IIA filaments drive cellular contractile system formation during interphase and mitosis. Mol. Biol. Cell 27, 1465–1478.

Fenix, A.M., Neininger, A.C., Taneja, N., Hyde, K., Visetsouk, M.R., Garde, R.J., Liu, B., Nixon, B.R., Manalo, A.E., Becker, J.R., et al. (2018). Muscle-specific stress fibers give rise to sarcomeres in cardiomyocytes. Elife 7.

Geiger, B., and Yamada, K.M. (2011). Molecular architecture and function of matrix adhesions. Cold Spring Harb. Perspect. Biol. 3, a005033..

Graessl, M., Koch, J., Calderon, A., Kamps, D., Banerjee, S., Mazel, T., Schulze, N., Jungkurth, J.K., Patwardhan, R., Solouk, D., et al. (2017). An excitable Rho GTPase signaling network generates dynamic subcellular contraction patterns. 216, 4271–4285.

Hayakawa, K., Tatsumi, H., and Sokabe, M. (2011). Actin filaments function as a tension sensor by tension-dependent binding of cofilin to the filament. J. Cell Biol. 195, 721–727.

Henson, J.H., Ditzler, C.E., Germain, A., Irwin, P.M., Vogt, E.T., Yang, S., Wu, X., and Shuster, C.B. (2017). The ultrastructural organization of actin and myosin II filaments in the contractile ring: New support for an old model of cytokinesis. Mol. Biol. Cell 28, 613–623.

Hotulainen, P., and Lappalainen, P. (2006). Stress fibers are generated by two distinct actin assembly mechanisms in motile cells. J. Cell Biol. 173, 383–394.

Hu, S., Dasbiswas, K., Guo, Z., Tee, Y.-H., Thiagarajan, V., Hersen, P., Chew, T.-L., Safran, S.A., Zaidel-Bar, R., and Bershadsky, A.D. (2017). Long-range self-organization of cytoskeletal myosin II filament stacks. Nat. Cell Biol. 19, 133–141.

Jacobelli, J., Bennett, F.C., Pandurangi, P., Tooley, A.J., and Krummel, M.F. (2009). Myosin-IIA and ICAM-1 Regulate the Interchange between Two Distinct Modes of T Cell Migration. J. Immunol. 182, 2041–2050.

Jiu, Y., Kumari, R., Fenix, A.M., Schaible, N., Liu, X., Varjosalo, M., Krishnan, R., Burnette, D.T., and Lappalainen, P. (2019). Myosin-18B Promotes the Assembly of Myosin II Stacks for Maturation of Contractile Actomyosin Bundles. Curr. Biol. 29, 81-92.e5.

Kassianidou, E., Brand, C.A., Schwarz, U.S., and Kumar, S. (2017). Geometry and network connectivity govern the mechanics of stress fibers. Proc. Natl. Acad. Sci. U. S. A. 114, 2622–2627.

Kim, H.Y., and Davidson, L.A. (2011). Punctuated actin contractions during convergent extension and their permissive regulation by the non-canonical Wnt-signaling pathway. J. Cell Sci. 124, 635–646.

Kim, D.-H., Cho, S., and Wirtz, D. (2014). Tight coupling between nucleus and cell migration through the perinuclear actin cap. J. Cell Sci. 127, 2528–2541.

Kim, D.H., Khatau, S.B., Feng, Y., Walcott, S., Sun, S.X., Longmore, G.D., and Wirtz, D. (2012). Actin cap associated focal adhesions and their distinct role in cellular mechanosensing. Sci. Rep. 2, 1–13.

Kraus, F., Miron, E., Demmerle, J., Chitiashvili, T., Budco, A., Alle, Q., Matsuda, A., Leonhardt, H., Schermelleh, L., and Markaki, Y. (2017). Quantitative 3D structured illumination microscopy of nuclear structures. Nat. Protoc. 12, 1011–1028.

Krishnan, R., Park, C.Y., Lin, Y.-C., Mead, J., Jaspers, R.T., Trepat, X., Lenormand, G., Tambe, D., Smolensky, A. V., Knoll, A.H., et al. (2009). Reinforcement versus Fluidization in Cytoskeletal Mechanoresponsiveness. PLoS One 4, e5486.

Kumari, R., Jiu, Y., Carman, P.J., Tojkander, S., Kogan, K., Varjosalo, M., Gunning, P.W., Dominguez, R., and Lappalainen, P. (2020). Tropomodulins Control the Balance between Protrusive and Contractile Structures by Stabilizing Actin-Tropomyosin Filaments. Curr. Biol. 30, 767-778.e5.

Laplante, C., Huang, F., Tebbs, I.R., Bewersdorf, J., and Pollard, T.D. (2016). Molecular organization of cytokinesis nodes and contractile rings by super-resolution fluorescence microscopy of live fission yeast. Proc. Natl. Acad. Sci. U. S. A. 113, E5876–E5885.

Lee, S., and Kumar, S. (2020). Cofilin is required for polarization of tension in stress fiber networks during migration. J. Cell Sci. jcs.243873.

Lehtimäki, J., Hakala, M., and Lappalainen, P. (2016). Actin Filament Structures in Migrating Cells. In Handbook of Experimental Pharmacology, pp. 1–30.

Lehtimäki, J.I., Fenix, A.M., Kotila, T.M., Balistreri, G., Paavolainen, L., Varjosalo, M., Burnette, D.T., and Lappalainen, P. (2017). UNC-45a promotes myosin folding and stress fiber assembly. J. Cell Biol. jcb.201703107.

Livne, A., and Geiger, B. (2016). The inner workings of stress fibers - from contractile machinery to focal adhesions and back. J. Cell Sci. 129, 1293–1304.

Logue, J.S., Cartagena-Rivera, A.X., Baird, M.A., Davidson, M.W., Chadwick, R.S., and Waterman, C.M. (2015). Erk regulation of actin capping and bundling by Eps8 promotes cortex tension and leader bleb-based migration. Elife 4, 1–31.

Lu, Y., Zhang, Y., Pan, M.-H., Kim, N.-H., Sun, S.-C., and Cui, X.-S. (2017). Daam1 regulates fascin for actin assembly in mouse oocyte meiosis. Cell Cycle 16, 1350–1356.

Luxton, G.W.G., Gomes, E.R., Folker, E.S., Vintinner, E., and Gundersen, G.G. (2010). Linear arrays of nuclear envelope proteins harness retrograde actin flow for nuclear movement. Science 329, 956–959.

Mason, F.M., Tworoger, M., and Martin, A.C. (2013). Apical domain polarization localizes actin– myosin activity to drive ratchet-like apical constriction. Nat. Cell Biol. 15, 926–936.

Michaux, J.B., Robin, F.B., McFadden, W.M., and Munro, E.M. (2018). Excitable RhoA dynamics drive pulsed contractions in the early C. elegans embryo. J. Cell Biol. 217, 4230–4252.

Munjal, A., Philippe, J.-M., Munro, E., and Lecuit, T. (2015). A self-organized biomechanical network drives shape changes during tissue morphogenesis. Nature 524, 351–355.

Munro, E., Nance, J., and Priess, J.R. (2004). Cortical flows powered by asymmetrical contraction transport PAR proteins to establish and maintain anterior-posterior polarity in the early C. elegans embryo. Dev. Cell 7, 413–424.

Naumanen, P., Lappalainen, P., and Hotulainen, P. (2008). Mechanisms of actin stress fibre assembly. J. Microsc. 231, 446–454.

Petrie, R.J., Koo, H., and Yamada, K.M. (2014). Generation of compartmentalized pressure by a nuclear piston governs cell motility in a 3D matrix. Science (80-.). 345, 1062–1065.

Prager-Khoutorsky, M., Lichtenstein, A., Krishnan, R., Rajendran, K., Mayo, A., Kam, Z., Geiger, B., and Bershadsky, A.D. (2011). Fibroblast polarization is a matrix-rigidity-dependent process controlled by focal adhesion mechanosensing. Nat. Cell Biol. 13, 1457–1465.

Rajakylä, E.K., Lehtimäki, J.I., Acheva, A., Schaible, N., Lappalainen, P., Krishnan, R., and Tojkander, S. (2020). Assembly of Peripheral Actomyosin Bundles in Epithelial Cells Is Dependent on the CaMKK2/AMPK Pathway. Cell Rep. 30, 4266-4280.e4.

Rauzi, M., Lenne, P.F., and Lecuit, T. (2010). Planar polarized actomyosin contractile flows control epithelial junction remodelling. Nature 468, 1110–1115.

Ruprecht, V., Wieser, S., Callan-Jones, A., Smutny, M., Morita, H., Sako, K., Barone, V., Ritsch-Marte, M., Sixt, M., Voituriez, R., et al. (2015). Cortical contractility triggers a stochastic switch to fast amoeboid cell motility. Cell 160, 673–685.

Schindelin, J., Arganda-Carreras, I., Frise, E., Kaynig, V., Longair, M., Pietzsch, T., Preibisch, S., Rueden, C., Saalfeld, S., Schmid, B., et al. (2012). Fiji: An open-source platform for biological-image analysis. Nat. Methods 9, 676–682.

Schulze, N., Graessl, M., Blancke Soares, A., Geyer, M., Dehmelt, L., and Nalbant, P. (2014). FHOD1 regulates stress fiber organization by controlling the dynamics of transverse arcs and dorsal fibers. J. Cell Sci. 127, 1379–1393.

Shcherbakova, D.M., Baloban, M., Emelyanov, A. V., Brenowitz, M., Guo, P., and Verkhusha, V. V. (2016). Bright monomeric near-infrared fluorescent proteins as tags and biosensors for multiscale imaging. Nat. Commun. 7, 1–12.

Shcherbo, D., Murphy, C.S., Ermakova, G. V., Solovieva, E.A., Chepurnykh, T. V., Shcheglov, A.S., Verkhusha, V. V., Pletnev, V.Z., Hazelwood, K.L., Roche, P.M., et al. (2009). Far-red fluorescent tags for protein imaging in living tissues. Biochem. J. 418, 567–574.

Small, J.V., Rottner, K., Kaverina, I., and Anderson, K.I. (1998). Assembling an actin cytoskeleton for cell attachment and movement. Biochim. Biophys. Acta - Mol. Cell Res. 1404, 271–281.

Solon, J., Kaya-Çopur, A., Colombelli, J., and Brunner, D. (2009). Pulsed Forces Timed by a Ratchet-like Mechanism Drive Directed Tissue Movement during Dorsal Closure. Cell 137, 1331–1342.

Svitkina, T.M. (2020). Actin Cell Cortex: Structure and Molecular Organization. Trends Cell Biol. 30, 556–565.

Tee, Y.H., Shemesh, T., Thiagarajan, V., Hariadi, R.F., Anderson, K.L., Page, C., Volkmann, N., Hanein, D., Sivaramakrishnan, S., Kozlov, M.M., et al. (2015). Cellular chirality arising from the self-organization of the actin cytoskeleton. Nat. Cell Biol. 17, 445–457.

Thomas, D.G., Yenepalli, A., Denais, C.M., Rape, A., Beach, J.R., Wang, Y., Schiemann, W.P., Baskaran, H., Lammerding, J., and Egelhoff, T.T. (2015). Non-muscle myosin IIB is critical for nuclear translocation during 3D invasion. J. Cell Biol. 210, 583–594.

Tojkander, S., Gateva, G., Schevzov, G., Hotulainen, P., Naumanen, P., Martin, C., Gunning, P.W., and Lappalainen, P. (2011). A molecular pathway for myosin II recruitment to stress fibers. Curr. Biol. 21, 539–550.

Tojkander, S., Gateva, G., and Lappalainen, P. (2012). Actin stress fibers - assembly, dynamics and biological roles. J. Cell Sci. 125, 1855–1864.

Tojkander, S., Gateva, G., Husain, A., Krishnan, R., and Lappalainen, P. (2015). Generation of contractile actomyosin bundles depends on mechanosensitive actin filament assembly and disassembly. Elife 4, e06126.

Tojkander, S., Ciuba, K., and Lappalainen, P. (2018). CaMKK2 Regulates Mechanosensitive Assembly of Contractile Actin Stress Fibers. Cell Rep. 24, 11–19.

Tolić-Nørrelykke, I.M., Butler, J.P., Chen, J., and Wang, N. (2002). Spatial and temporal traction response in human airway smooth muscle cells. Am. J. Physiol. Physiol. 283, C1254–C1266.

Vasquez, C.G., Tworoger, M., and Martin, A.C. (2014). Dynamic myosin phosphorylation regulates contractile pulses and tissue integrity during epithelial morphogenesis. J. Cell Biol. 206, 435–450.

Vicente-Manzanares, M., Ma, X., Adelstein, R.S., and Horwitz, A.R. (2009). Non-muscle myosin II takes centre stage in cell adhesion and migration. Nat. Rev. Mol. Cell Biol. 10, 778–790.

Wu, J., Kent, I.A., Shekhar, N., Chancellor, T.J., Mendonca, A., Dickinson, R.B., and Lele, T.P. (2014). Actomyosin pulls to advance the nucleus in a migrating tissue cell. Biophys. J. 106, 7–15.

Xu, K., Babcock, H.P., and Zhuang, X. (2012). Dual-objective STORM reveals three-dimensional filament organization in the actin cytoskeleton. Nat. Methods 9, 185–188.

Yamada, K.M., and Sixt, M. (2019). Mechanisms of 3D cell migration. Nat. Rev. Mol. Cell Biol. 20, 738–752.

Yamada, S., and Nelson, W.J. (2007). Localized zones of Rho and Rac activities drive initiation and expansion of epithelial cell-cell adhesion. J. Cell Biol. 178, 517–527.

Young, L.E., and Higgs, H.N. (2018). Focal Adhesions Undergo Longitudinal Splitting into Fixed-Width Units. Curr. Biol. 28, 2033-2045.e5.

Zaidel-Bar, R., Milo, R., Kam, Z., and Geiger, B. (2007). A paxillin tyrosine phosphorylation switch regulates the assembly and form of cell-matrix adhesions. J. Cell Sci. 120, 137–148.

