## Supplemental Figures S1-S4 and Movie Legends for "Generation of stress fibers through myosin-driven re-organization of the actin cortex"

### Supplemental items

**Supplementary Figures S1-S4**

**Supplementary Movie Legends 1-8**

### Generation of stress fibers through myosin-driven re-organization of the actin cortex

**Lehtimäki JI[1], Rajakylä EK[2], Tojkander S[2], Lappalainen P[1,*]**

1) Institute of Biotechnology, HiLIFE Institute of Biotechnology, P.O. Box 56, University of Helsinki, 00014, Helsinki, Finland
2) Section of Pathology, Department of Veterinary Biosciences, Faculty of Veterinary Medicine, Agnes Sjöbergin katu 2, University of Helsinki, 00014, Helsinki, Finland

***Corresponding author & lead contact**
Pekka Lappalainen, HiLIFE Institute of Biotechnology, P.O. Box 56, University of Helsinki, 00014, Helsinki, Finland


### Figure S1

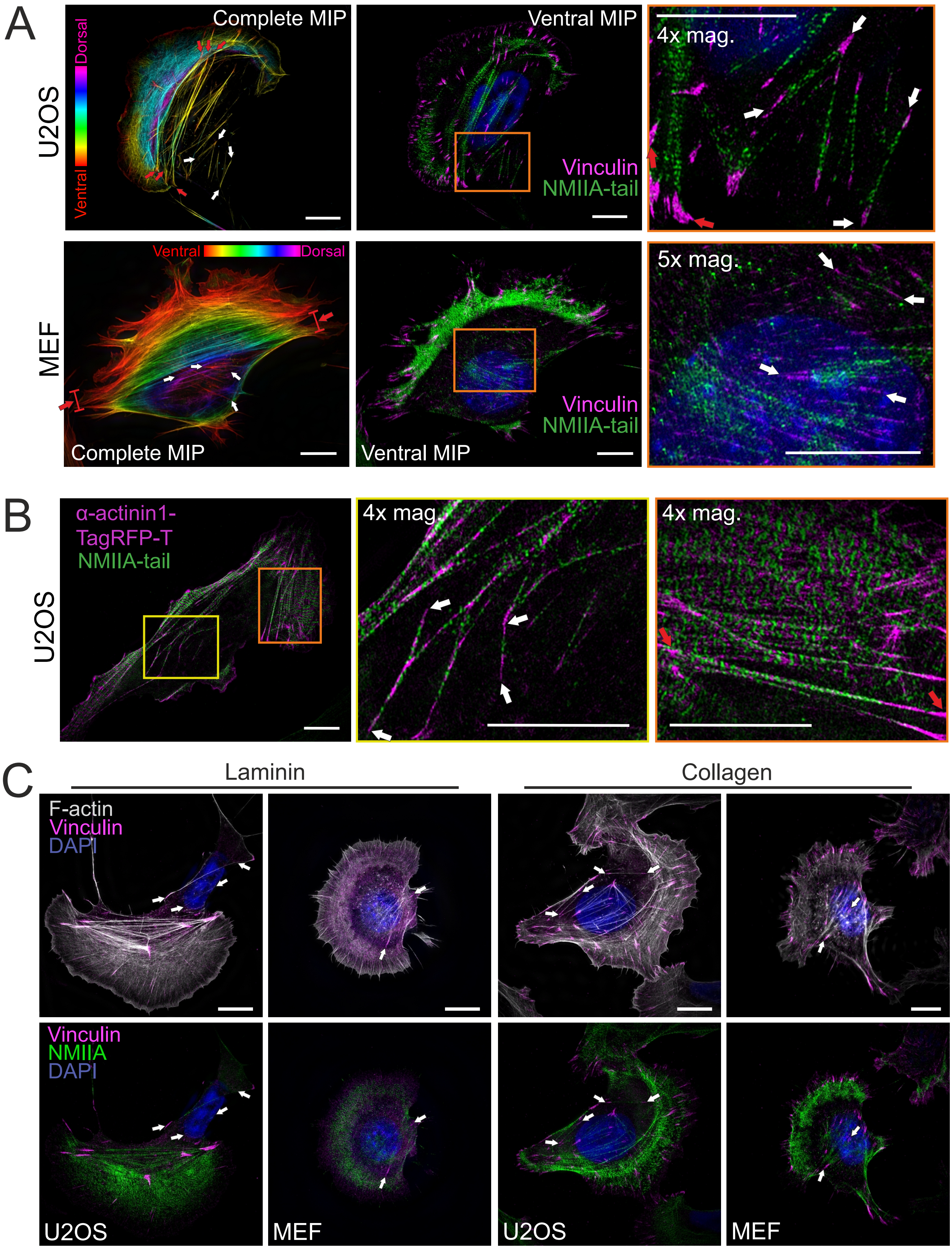
A
Complete MIP
Dorsal
Ventral
U2OS
Ventral MIP
Vinculin
NMIIA-tail
4x mag.
Ventral
Dorsal
MEF
Complete MIP
Ventral MIP
Vinculin
NMIIA-tail
5x mag.
B
α-actinin1-
TagRFP-T
NMIIA-tail
U2OS
4x mag.
4x mag.
C
Laminin
Collagen
F-actin
Vinculin
DAPI
Vinculin
NMIIA
DAPI
U2OS
MEF
U2OS
MEF

**Figure S1. Cortical stress fibers display periodic NMII - α-actinin pattern and assemble on different ECMs. A)** U20S and mouse embryonic fibroblast (MEF) cells (related to Fig. 1A), where vinculin and NMII were detected by specific antibodies, and F-actin and nucleus by fluorescent phalloidin and DAPI, respectively. Two panels on the left show temporal color-coded projection from the phalloidin staining, where red arrows highlight ventral stress fibers. At the middle, ventral plane of the cells show the localizations of vinculin and NMIIA tail. On the right, magnification of the boxed region form the middle panel demonstrating the presence of thin cortical stress fibers (white arrows) at the back of the cell and in the vicinity of the nucleus. **B)** α-actinin1 localization in ventral stress fibers and cortical stress fibers in a U2OS cell. 3D-SIM MIP with 4x magnifications of ventral stress fibers (orange box) display the periodic localization of the α–actinin and NMII in the ventral stress fibers (red arrows), whereas cortical stress fibers (4x mag., yellow box) display less regular pattern of α-actinin. **C)** 3D-SIM MIPs of U2OS and MEF cells cultured either on laminin or collagen coated dishes. F-actin was visualized by fluorescent phalloidin, focal adhesions by vinculin antibody, and nuclei by DAPI. Endogenous NMIIA tail domain was visualized by specific antibody. White arrows indicate examples of cortical stress fibers. Scalebars 10 μm and 5 μm for whole cell images and magnified areas, respectively.

### Figure S2

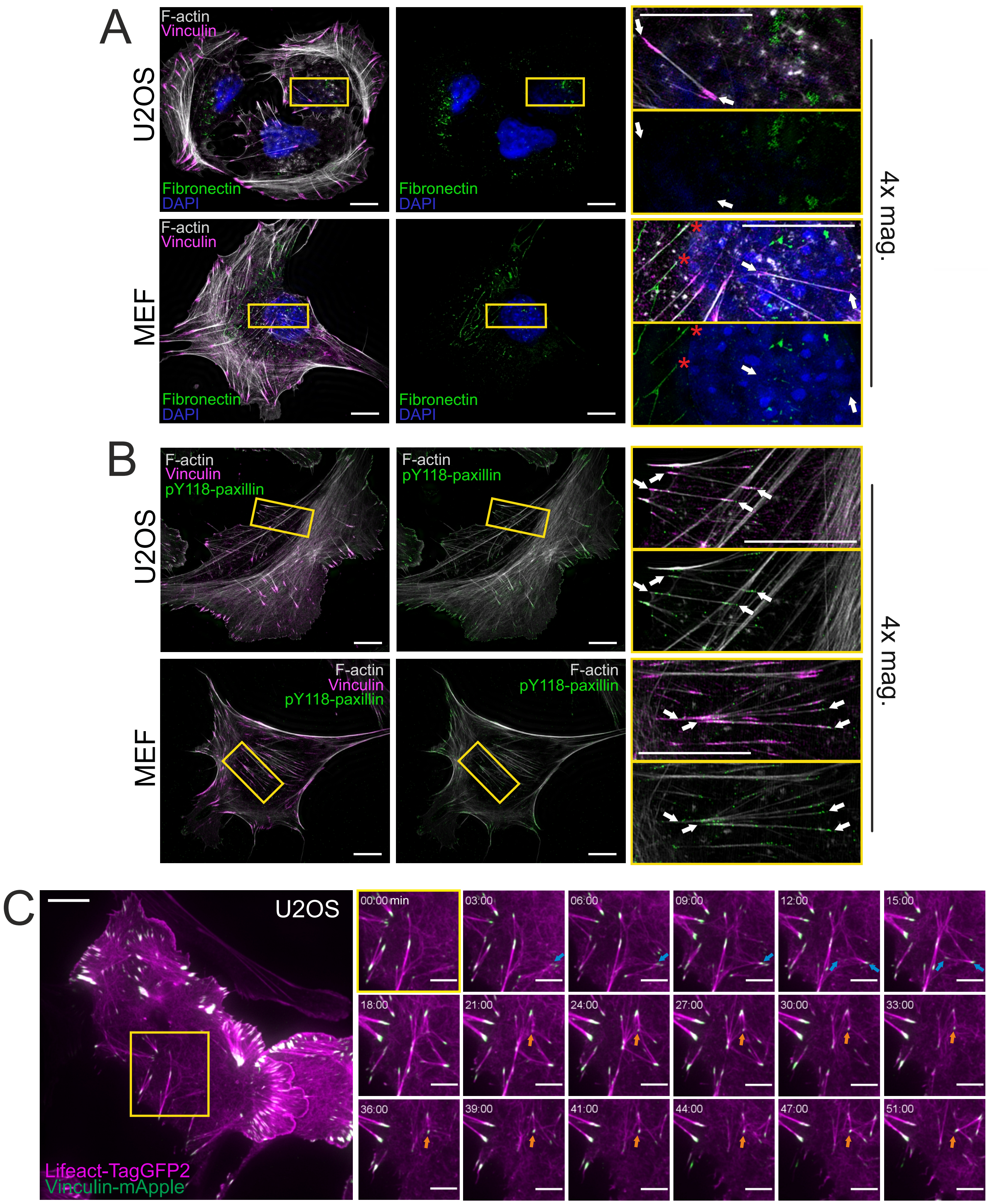
A
U2OS
F-actin
Vinculin
Fibronectin
DAPI
Fibronectin
DAPI
MEF
F-actin
Vinculin
Fibronectin
DAPI
Fibronectin
DAPI
4x mag.
B
U2OS
F-actin
Vinculin
pY118-paxillin
F-actin
pY118-paxillin
MEF
F-actin
Vinculin
pY118-paxillin
F-actin
pY118-paxillin
4x mag.
C
U2OS
Lifeact-TagGFP2
Vinculin-mApple
00:00 min
03:00
06:00
09:00
12:00
15:00
18:00
21:00
24:00
27:00
30:00
33:00
36:00
39:00
41:00
44:00
47:00
51:00

**Figure S2. Cortical stress fibers are not linked to fibrillar adhesions. A)** 3D-SIM MIPs of U2OS cells and MEFs cultured on uncoated high precision glass coverslips. Antibody detecting fibronectin was applied to visualize fibronectin deposits under the cells. 4x magnifications on the right (from yellow boxes) display cortical stress fibers in both cell types, and demonstrate that in U2OS cells they do not co-localize with the fibronectin (white arrows). For MEFs, some cortical stress fibers were associated with the fibronectin deposits (red asterisks). Nuclei were detected by DAPI. **B)** Antibody detecting the Y118-phosphorylated version of the paxillin was applied to discern the focal adhesions from fibrillar adhesions, which were reported to be devoid of paxillin phosphorylation (Zaidel-Bar et al., 2007). 3D-SIM MIPs from both U2OS and MEF cells on fibronectin-coated high precision coverslips were stained with phalloidin (F-actin) and an antibody to detect vinculin. 4x magnification of marked areas (yellow boxes) illustrate p-paxillin co-localization with vinculin at the focal adhesions in the ends of cortical stress fibers (white arrows). **C)** Cortical stress fiber assembly in a migrating U2OS cell expressing LifeAct-TagGFP2 (F-actin, magenta) and vinculin-mApple (focal adhesions, green) imaged by TIRF microscopy. Blue arrows exemplify the *de novo* generation of a cortical stress fiber, and the orange arrow highlights a vinculin positive adhesion that was used by several cortical stress fibers over the time. Imaging interval 30s. See also **Movie S2**. Scale bars 10 µm and 5 µm for the whole cell images and magnified images, respectively.

### Figure S3

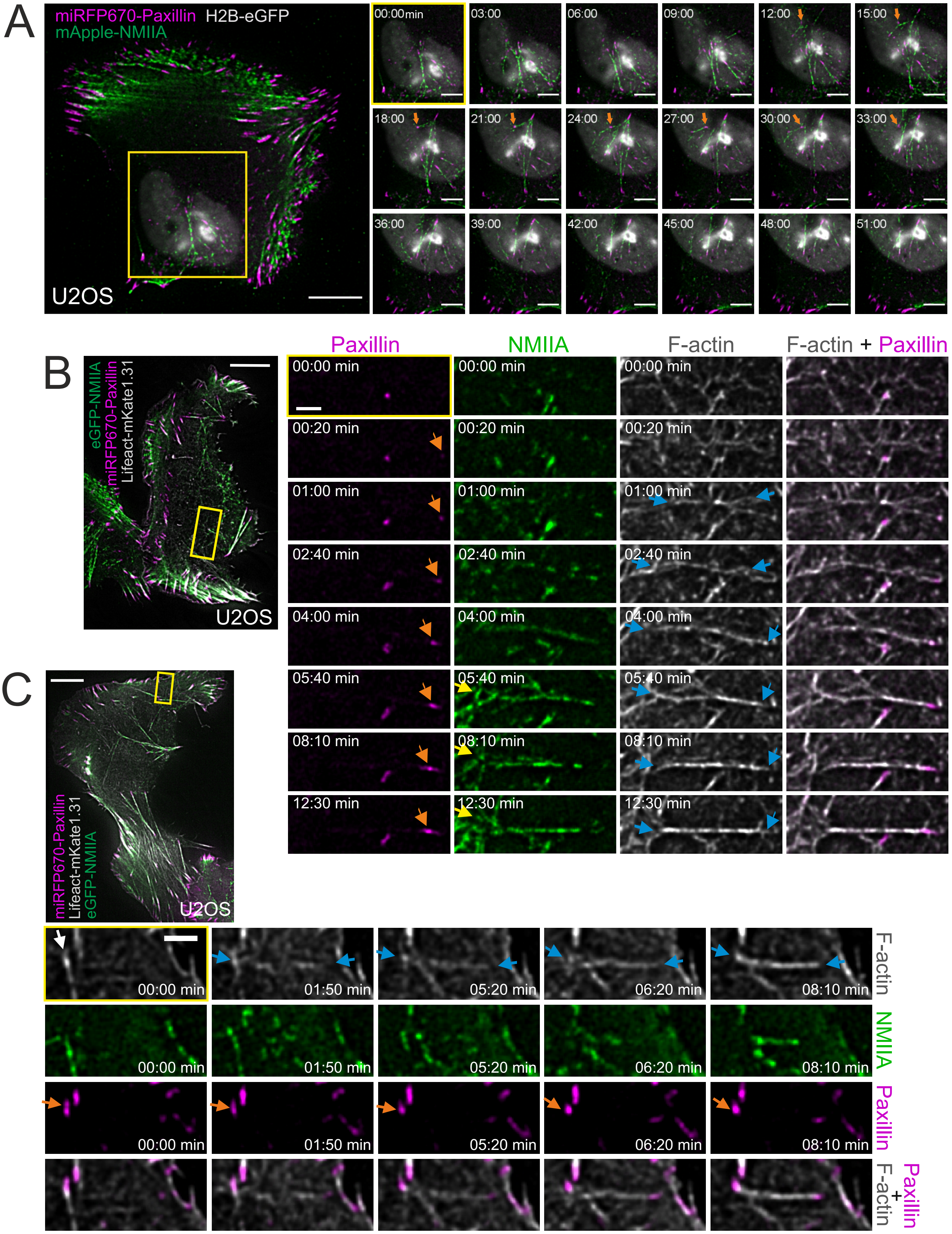
A
miRFP670-Paxillin H2B-eGFP
mApple-NMIIA
U2OS
00:00min
03:00
06:00
09:00
12:00
15:00
18:00
21:00
24:00
27:00
30:00
33:00
36:00
39:00
42:00
45:00
48:00
51:00
B
eGFP-NMIIA
miRFP670-Paxillin
Lifeact-mKate1.31
U2OS
Paxillin
NMIIA
F-actin
F-actin + Paxillin
00:00 min
00:20 min
01:00 min
02:40 min
04:00 min
05:40 min
08:10 min
12:30 min
C
miRFP670-Paxillin
Lifeact-mKate1.31
eGFP-NMIIA
U2OS
00:00 min
01:50 min
05:20 min
06:20 min
08:10 min
F-actin
NMIIA
Paxillin
Paxillin + F-actin

**Figure S3. Different ways to assemble cortical stress fibers from the ventral actomyosin cortex**. **A)** TIRF time-lapse imaging of a migrating a U2OS cell expressing histone H2B-eGFP (grey), mApple-NMIIA (green) and miRFP670-paxillin (focal adhesions, magenta). Selected time-lapse frames from the magnified area (yellow box) show examples of myosin pulses that assemble into cortical stress fibers underneath the nucleus, with concurrent maturation of paxillin-positive focal adhesions. Orange arrow illustrates how a single focal adhesion is used by two different cortical stress fibers assembling over the time (original imaging interval 30 s). Scale bars 10 μm and 5 μm for the whole cell image and magnified time-lapse frames, respectively. See also **Movie S3**. **B)** Example of a formation of cortical stress fiber in a migrating U2OS cell, where the one end of the stress fiber connect to a focal adhesion and the other one to a cortical actomyosin patch. Blue arrows in the selected, magnified time-lapse frames (from the yellow-boxed area) illustrate F-actin bundling, orange arrows highlight the maturing paxillin-positive focal adhesion (magenta, miRFP670-paxillin) and yellow arrows highlight the NMII patch where the stress fiber is connected from its other end. NMIIA (green) and F-actin (grey) were visualized by expressing eGFP-NMIIA and LifeAct-mKate1.31, respectively. **C)** Example of a focal adhesion exchanging from one cortical stress fiber to another one in a migrating U2OS cell expressing LifeAct-TagGFP2 (F-actin, grey), eGFP-NM-IIA (green) and miRFP670-paxillin (magenta). Selected time-lapse frames (magnified from the yellow boxed area) show that after disassembly of the preceding cortical stress fiber (white arrow), the newly forming cortical stress fiber (illustrated with blue arrows) engages itself to the pre-existing paxillin-positive focal adhesion (orange arrow). See also **Movie S5**. Scale bars 10 μm and 2 μm for whole cell images and magnified time-lapse frames, respectively. Original imaging interval 10s.

### Figure S4

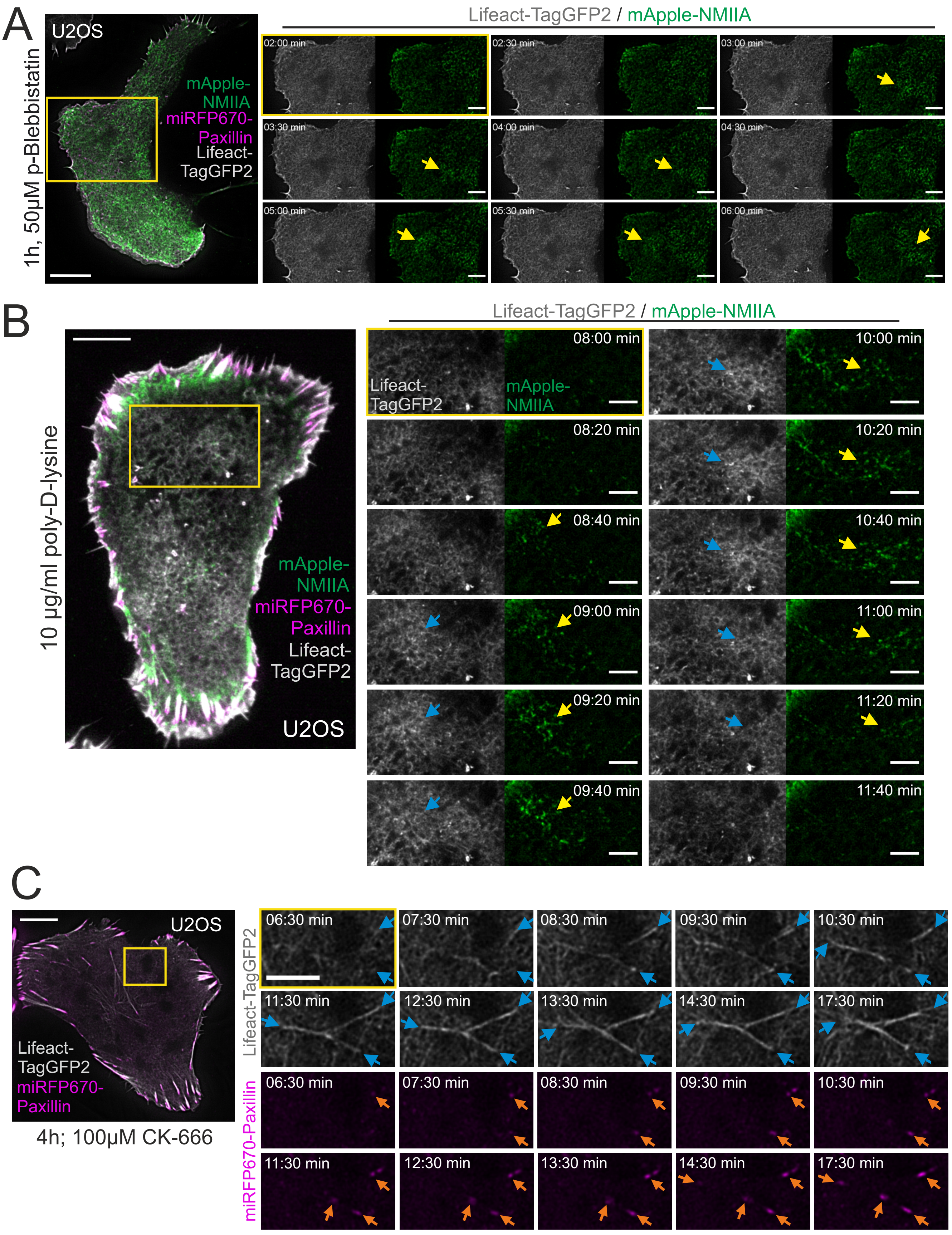
A
1h, 50μM p-Blebbistatin
U2OS
mApple-NMIIA
miRFP670-Paxillin
Lifeact-TagGFP2
Lifeact-TagGFP2 / mApple-NMIIA
02:00 min
02:30 min
03:00 min
03:30 min
04:00 min
04:30 min
05:00 min
05:30 min
06:00 min
B
10 µg/ml poly-D-lysine
mApple-NMIIA
miRFP670-Paxillin
Lifeact-TagGFP2
U2OS
Lifeact-TagGFP2 / mApple-NMIIA
Lifeact-TagGFP2
mApple-NMIIA
08:00 min
08:20 min
08:40 min
09:00 min
09:20 min
09:40 min
10:00 min
10:20 min
10:40 min
11:00 min
11:20 min
11:40 min
C
U2OS
Lifeact-TagGFP2
miRFP670-Paxillin
4h; 100μM CK-666
Lifeact-TagGFP2
miRFP670-Paxillin
06:30 min
07:30 min
08:30 min
09:30 min
10:30 min
11:30 min
12:30 min
13:30 min
14:30 min
17:30 min

**Figure S4. Cortical stress fiber assembly is dependent on functional NMII and integrin-based adhesions. A)** TIRF time-lapse images of a U2OS cell expressing mApple-NMIIA (green), miRFP670-paxillin (magenta) and LifeAct-TagGFP2 (grey), after 1 h incubation with 50 μM para-amino Blebbistatin. Blebbistatin does not cease myosin pulses highlighted with yellow arrows (see also Baird et al., 2017), but abolishes cortical stress fiber formation. Magnified time-lapse frames (from yellow boxed area) and **Movie S6** display the NMIIA pulses that are unable to bundle F-actin. **B)** TIRF time-lapse imaging of a U2OS cells plated onto poly-D-lysine coated dishes 24h prior imaging. Only cells that displayed partial integrin-mediated adhesion (based on miRFP670-paxillin, magenta) from their edges were imaged. Exemplary time-lapse images from the magnified region (yellow box) illustrate NMIIA pulses (mApple-NMIIA, green) that are unable to stabilize F-actin bundles. Note the blue arrows pointing out the transient accumulation of cortical actin (LifeAct-TagGFP2, grey), coinciding with the NMIIA pulses (yellow arrows). See also **Movie S7**. **C)** Cortical stress fiber formation in a U2OS cell transfected with miRFP670-paxillin and LifeAct-TagGFP2, and subjected to CK-666 for 4 hours prior onset of TIRF-imaging. Selected time-lapse frames from the magnified regions show an example of cortical stress fiber formation (blue arrows) and reinforcement of focal adhesions (orange arrows). See also **Movie S8**. Scale bars 10 μm and 5 μm for the whole cell images and magnifications, respectively. Time-lapse imaging interval 30 s (A,C) and 20 s for (B), respectively.

#### Supplementary movie legends

**Movie S1. Cortical stress fiber formation at the ventral actin cortex of migrating U2OS and MEF cells.** U2OS (on the left) and MEF (on the right) cells expressing LifeAct-TagGFP2 (grey/magenta, respectively) and mApple-vinculin (magenta/green, respectively), migrating on a fibronectin (10 μg/ml)-coated high-precision imaging dish. Related to Figure 2A-B. Time-lapse ring-TIRFM videos were recorded with Deltavision OMX SR with 30 s imaging interval. Playback rate 15 frames/s; PNG compressed video. Scalebars 10 μm.

**Movie S2. NMIIA pulsation and cortical stress fiber assembly at the trailing edge of a migrating U2OS cells.** U2OS cells expressing either LifeAct-TagGFP2 (magenta) plus mApple-vinculin (green) (on the left) or mApple-vinculin eGFP-NMIIA (green) and mApple-vinculin (magenta) (on the right) migrating on a fibronectin (10 μg/ml)-coated high-precision imaging dishes. Related to Figure S2C and 3D, respectively. Time-lapse ring-TIRFM videos were recorded with Deltavision OMX SR with 30 s imaging interval. Playback rate 15 frames/s; PNG compressed video. Scalebars 10 μm.

**Movie S3. NMIIA pulses assemble into cortical stress fibers under the nucleus of migrating U2OS cells.** U2OS cells expressing either LifeAct-TagGFP2 (grey), histone-H2B-mCherry (blue) and miRFP670-paxillin (magenta), (on the left) or Histone H2B-eGFP (grey), mApple-NMIIA (green) and miRFP670-paxillin (magenta), (on the right), both migrating on a fibronectin (10 μg/ml)-coated high-precision imaging dish. Related to Figure 3A and S3A. Time-lapse ring-TIRFM videos were recorded with Deltavision OMX SR with 30 s and 20 s imaging intervals, respectively. Playback rate 20 frames/s; PNG compressed video. Scalebars 10 μm.

**Movies S4 and S5. NMIIA pulses coordinate the cortical stress fiber assembly in migrating U2OS cells.** U2OS cells expressing LifeAct-TagGFP2 (grey), mApple-NMIIA (green) and miRFP670-paxillin (magenta), migrating on a fibronectin (10 µg/ml)-coated high-precision imaging dish. Related to Figure 4A and S3B-C, respectively. Time-lapse ring-TIRFM videos were recorded with Deltavision OMX SR with 30 s and 10 s imaging intervals (for S4 and S5, respectively). Playback rate 15 and 20 frames/s, respectively. PNG compressed video. Scalebars 10 µm.

**Movie S6. Inhibiting NMII activity abolishes cortical stress fiber assembly.** U2OS cell expressing LifeAct-TagGFP2 (grey), mApple-NMIIA (green) and miRFP670-paxillin (magenta) migrating on a fibronectin (10 µg/ml) -coated high precision imaging dish after one-hour treatment with 50 µM p-amino-Blebbistatin. Related to Figure S4A. Time-lapse ring-TIRFM video was recorded with Deltavision OMX SR with 30 s imaging interval. Playback rate 15 frames/s; PNG compressed video. Scalebar 10 µm.

**Movie S7. Inhibiting integrin-based ECM adhesion obstructs cortical stress fiber formation** U2OS cell expressing LifeAct-TagGFP2 (grey), mApple-NMIIA (green) and miRFP670-paxillin (magenta) on a poly-D-lysine (10 µg/ml)-coated high-precision imaging dish. Imaging started 24 h post plating. Related to Figure S4B. Time-lapse ring-TIRFM video was recorded with Deltavision OMX SR with 20 s imaging interval. Playback rate 20 frames/s; PNG compressed video. Scalebar 10 µm.

**Movie S8. CK-666 mediated Arp2/3 inhibition does not prevent cortical stress fiber assembly.** U2OS cell expressing LifeAct-TagGFP2 (grey) and miRFP670-paxillin (magenta) migrating on a fibronectin (10 µg/ml)-coated high-precision imaging dish after 4-hour treatment with 100 µM CK-666. Related to Figure S4C. Time-lapse ring-TIRFM video was recorded with Deltavision OMX SR with 30s imaging interval. Playback rate 15 frames/s; PNG compressed video. Scalebar 10 µm.

### References

Baird, M.A., Billington, N., Wang, A., Adelstein, R.S., Sellers, J.R., Fischer, R.S., and Waterman, C.M. (2017). Local pulsatile contractions are an intrinsic property of the myosin 2A motor in the cortical cytoskeleton of adherent cells. Mol. Biol. Cell *28*, 240–251.

Zaidel-Bar, R., Milo, R., Kam, Z., and Geiger, B. (2007). A paxillin tyrosine phosphorylation switch regulates the assembly and form of cell-matrix adhesions. J. Cell Sci. *120*, 137–148.
